# mRNA Degradation Rates Are Coupled to Metabolic Status in Mycobacteria

**DOI:** 10.1101/595199

**Authors:** Diego A. Vargas-Blanco, Ying Zhou, Luis Gutierrez Zamalloa, Tim Antonelli, Scarlet S. Shell

## Abstract

The success of *Mycobacterium tuberculosis* (Mtb) as a human pathogen is due in part to its ability to survive stress conditions, such as hypoxia or nutrient deprivation, by entering non-growing states. In these low-metabolic states, Mtb can tolerate antibiotics and develop genetically encoded antibiotic resistance, making its metabolic adaptation to stress crucial for survival. Numerous bacteria, including Mtb, have been shown to reduce their rates of mRNA degradation under growth limitation and stress. While the existence of this response appears to be conserved across species, the underlying bacterial mRNA stabilization mechanisms remains unknown. To better understand the biology of non-growing mycobacteria, we sought to identify the mechanisms by which mRNA stabilization occurs using the non-pathogenic model *Mycobacterium smegmatis*. We found that mRNA half-life was responsive to energy stress, with carbon starvation and hypoxia causing global mRNA stabilization. This global mRNA stabilization was rapidly reversed when hypoxia-adapted cultures were re-exposed to oxygen, even in the absence of new transcription. The stringent response and RNase protein levels did not explain mRNA stabilization, nor did transcript abundance. This led us to hypothesize that metabolic changes during growth cessation impact the activity of degradation proteins, increasing mRNA stability. Indeed, bedaquiline and isoniazid, two drugs with opposing effects on cellular energy status, had opposite effects on mRNA half-lives in growth-arrested cells. Taken together, our results indicate that mRNA stability in mycobacteria is not directly regulated by growth status, but rather seems to be dependent on the status of energy metabolism.

**IMPORTANCE:** The logistics of treating tuberculosis are difficult, requiring multiple drugs for at least six months. Mtb is able to survive within the human host in part by entering non-growing states in which it is metabolically less active, thus rendering it less susceptible to antibiotics. Basic knowledge on how Mtb survives during these low-metabolic states is incomplete, and we postulate that optimized energy resource management –such as transcriptome stabilization—is important for survival. Here we report that mRNA stabilization (increased mRNA half-lives) is a common feature of mycobacteria under stress (e.g. hypoxia and nutrient deprivation) but is not dependent on the mechanisms that have been most often postulated in the literature. Finally, we found that mRNA stability and growth status can be decoupled by a drug that causes growth arrest but increases metabolic activity, indicating that mRNA stability responds to metabolic status rather than to growth rate changes per se. Our findings suggest a need to re-orient the study of global mRNA stabilization to identify novel mechanisms that are presumably responsible.

## INTRODUCTION

Most bacteria periodically face environmental conditions that are unfavorable for growth. To overcome such challenges, bacteria must adapt both their gene expression profiles and their energy usage. Regulation of mRNA turnover can contribute to both of these. However, the mechanisms by which mRNA turnover is carried out and regulated remain poorly understood, particularly in mycobacteria.

During infection, the human pathogen *Mycobacterium tuberculosis* (Mtb) faces not only the immune response and antibiotics, but also multiple non-optimal microenvironments, such as hypoxia and nutrient starvation within the granuloma (1, 2). Regulation of mRNA turnover appears to contribute to adaptation to such conditions. A global study of mRNA decay in Mtb showed a dramatic increase in transcriptome stability—measured as increased mRNA half-lives— in response to hypoxia, when compared to log phase growth in oxygen-rich conditions (3). This suggests that mRNA stabilization is important for energy conservation in the energy-limited environments that Mtb encounters during infection. Similar responses have been shown for other bacteria under stress conditions that slow or halt growth, including carbon deprivation, stationary phase, and temperature shock (4–13). However, the mechanisms responsible for global regulation of mRNA stability in prokaryotes have yet to be elucidated.

In better studied bacteria such as *E. coli* and *B. subtilis,* the major ribonucleases (RNases) involved in mRNA processing and decay are RNase E and RNase Y, respectively. A conventional model for RNA decay in *E. coli* start with an endonucleolytic cleavage event usually carried by RNase E in AU-rich regions, particularly in mRNA substrates that possess a 5’ monophosphate (14–16). The resulting 5’ monophosphorylated fragments are rapidly cleaved by RNase E, resulting in shorter fragments that can be fully degraded by exonucleases such as PNPase, RNase II, and RNase R (17, 18). mRNA degradation seems to be coordinated by formation of a complex known as the degradosome. In *E. coli,* RNase E serves as the scaffold for this multiprotein complex that comprises RNA helicases, the glycolytic enzyme enolase, and PNPase (19–23). Other organisms that encode RNase E form similar degradosomes (24, 25). In organisms where RNase E is not present, RNase Y and/or RNase J seem to assume the scaffold function (26–28). Mycobacteria encode RNase E, but efforts to define the mycobacterial degradosome have produced inconsistent results (29, 30). It is unclear if degradosome reorganization or dissolution contribute to the global regulation of mRNA degradation under stress conditions in any bacteria. Interestingly, the importance of degradosome formation in *E. coli* varies depending on the carbon sources provided, suggesting specific links between RNase E degradosomes and metabolic capabilities (31). Furthermore, the chaperones DnaK and CsdA can become degradosome components in *E. coli* under certain stresses (20, 32, 33).

Global transcript stabilization in stressed bacteria could plausibly result from reduced RNase abundance, reduced RNase activity, and/or reduced accessibility of transcripts to degradation proteins. In *E. coli* it has been shown that multiple stressors can upregulate RNase R, possibly as a way to overcome ribosome misassembly (34, 35), and that RNase III levels decrease under cold-shock and stationary phase (36). Surprisingly, protein levels for most putative RNA degradation proteins in *M. tuberculosis* remain unaltered under hypoxic conditions (37), which suggests that mRNA degradation is not necessarily regulated at the level of RNase abundance in mycobacteria. However, there is evidence that RNase activity may be regulated. For example, proteins such as RraA and RraB can alter the function of the RNase E-based degradosome in *E. coli* (38). Translating ribosomes can mask mRNA cleavage sites, and, indeed, transcription-translation dissociation experiments showed that ribosome-free mRNAs were highly unstable (39). Furthermore, in some actinomycetes PNPase might be regulated by the stringent response alarmone guanosine tetraphosphate (ppGpp) (40, 41). In Gram-negative bacteria ppGpp is usually synthesized by RelA, which is activated in the presence of uncharged tRNAs, or by the ppGpp synthase/hydrolase SpoT during fatty acid starvation (42). In Gram-positive bacteria, ppGpp is commonly synthesized by a dual RelA/SpoT homolog (43–45). Diverse bacteria adapt to stress using ppGpp in different pathways, which generally result in halting the synthesis of stable RNA (tRNAs and rRNAs), while upregulating stress-associated genes and downregulating those associated with cell growth (45–50). Recent reports in two actinomycetes –*Streptomyces coelicolor* and *Nonomuraea*—showed that, at physiological levels, ppGpp inhibited the enzymatic activity of PNPase (40, 41), suggesting that the stringent response could directly stabilize mRNA as part of a broader response to energy starvation.

Another explanation for stress-induced transcript stabilization could be that reduced transcript abundance directly leads to increased transcript stability. mRNA abundance and half-life were reported to be inversely correlated in multiple bacteria including Mtb (3, 8, 51, 52), and mRNA abundance is lower on a per-cell basis for most transcripts in non-growing bacteria. In *Caulobacter crescentus*, subcellular localization of mRNA degradation proteins may play a role in global mRNA stability (53, 54). Nevertheless, the causal relationships between translation, mRNA abundance, RNase expression, and mRNA stability in non-growing bacteria remain largely untested.

Given the importance of adaptation to energy starvation for mycobacteria, we sought to investigate the mechanisms by which mRNA stability is globally regulated. Here we show that the global mRNA stabilization response occurs also in *Mycobacterium smegmatis*—a non-pathogenic model commonly used to study the basic biology of mycobacteria —under hypoxia and carbon starvation. We found that hypoxia-induced mRNA stability is rapidly reversible, with re-aeration causing immediate mRNA destabilization even in the absence of protein synthesis. As expected, we found that transcript levels from hypoxic cells are lower on a per-cell basis compared to those from aerated cultures. However, our data are inconsistent with a model in which mRNA abundance dictates degradation rate as has been shown for log-phase *E. coli* (51) and *Lactococcus lactis* (52). Instead, our findings support the idea that mRNA stability is rapidly tuned in response to alterations in energy metabolism. This effect does not require the stringent response or changes in abundance of RNA degradation proteins, and it can be decoupled from growth status.

## RESULTS

### mRNA is stabilized as a response to carbon starvation and hypoxic stress in *Mycobacterium smegmatis*

The mRNA pools of *E. coli* and other well-studied bacteria were reported to be globally stabilized during conditions of stress, resulting in increased mRNA half-lives (3–13). In 2013, Rustad *et al.* reported a similar phenomenon in Mtb under hypoxia and during cold shock (3). We sought to establish *M. smegmatis* as a model for study of the mechanistic basis of mRNA stabilization in mycobacteria under stress conditions. We therefore subjected *M. smegmatis* to hypoxic and carbon starvation stresses, and measured mRNA half-lives for a subset of genes by blocking transcription with rifampicin (RIF) and measuring mRNA abundance at multiple time points using quantitative PCR (qPCR). Indeed, we observed that all of the analyzed transcripts had increased half-lives under hypoxia when compared to log phase normoxic cultures and, similarly, transcripts were more stable in carbon starvation than in rich media (Fig. 1A and 1B). Thus, *M. smegmatis* appears to be a suitable model organism for investigating the mechanisms of stress-induced mRNA stabilization in mycobacteria. We used a variation of the Wayne model (55) to produce a gradual transition from aerated growth to hypoxia-induced growth arrest by sealing cultures in vials with defined headspace ratios and allowing them to slowly deplete the available oxygen (Fig. 1C). We noted that transcripts became progressively more stable as oxygen levels dropped and growth ceased; 40 hours after sealing the vials, mRNA half-lives were too long to reliably measure by our methodology. We sought to focus our studies on the mechanisms that underlie the initial mRNA stabilization process during the transition into hypoxia-induced growth arrest. We therefore conducted subsequent experiments 18 hours after sealing the vials, when growth had nearly ceased and transcripts were 9-fold to 25-fold more stable than during log phase growth. We hereafter refer to this condition as 18 h hypoxia.

**Figure 1.**
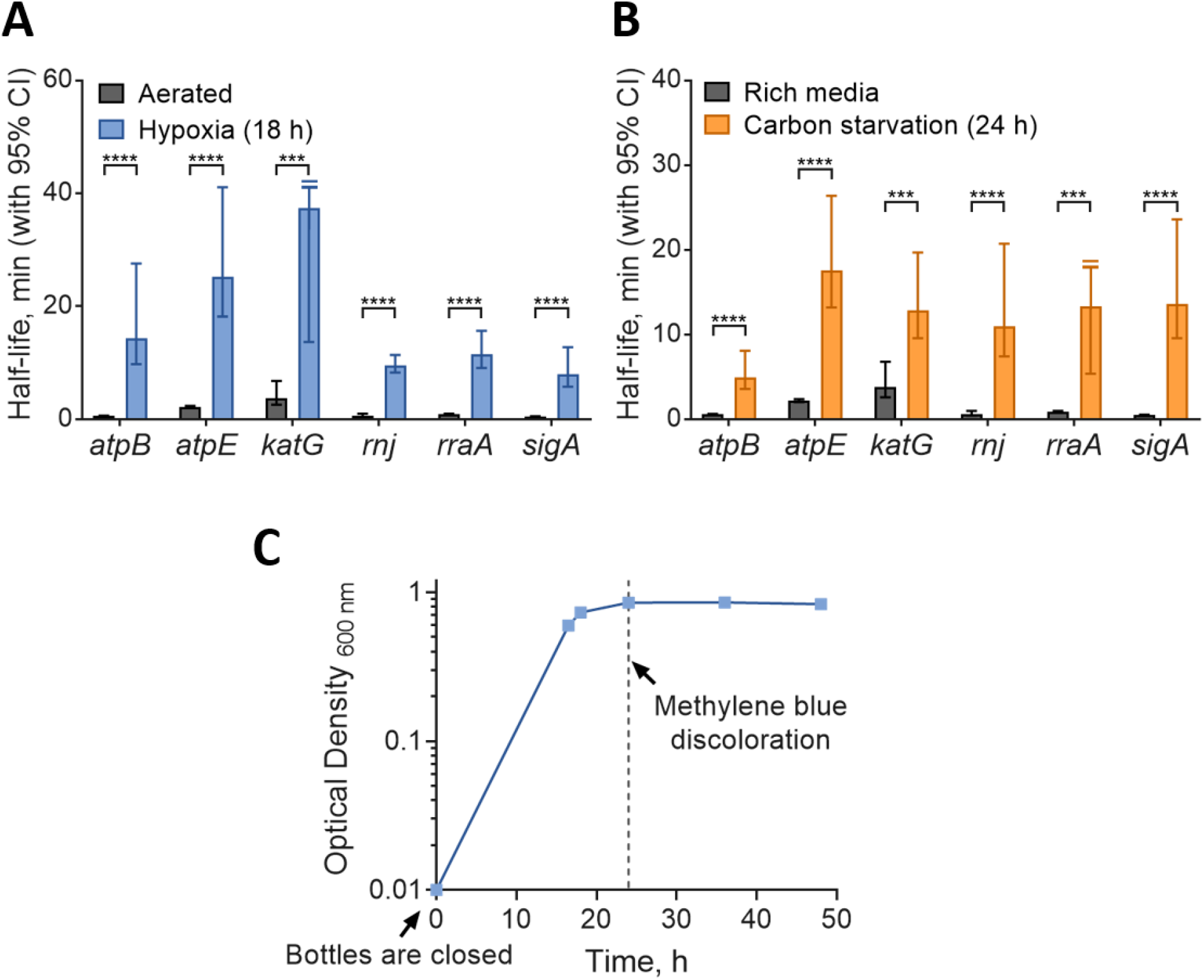
Transcript half-lives are increased in response to hypoxia and carbon starvation stress. Transcript half-lives for the indicated genes were measured for *M. smegmatis* mc^2^155 after blocking transcription with 150 μg·mL^−1^ RIF. RNA samples were collected (A) during log phase normoxia, and hypoxia (18 hours after closing the bottles); or (B) during log phase in 7H9 supplemented with ADC, glycerol, and Tween 80 (rich media) or 7H9 with Tyloxapol only (carbon starvation, 24 hours). Degradation rates were compared using linear regression (*n*=3), and half-lives were determined by the reciprocal of the best-fit slope. Error bars: 95% CI. *** *p*<0.001; **** *p*<0.0001. When a slope of zero was included in the 95% CI (indicating no degradation), the upper limit for half-life was unbounded, indicated by a clipped error bar with a double line. (C) Growth kinetics for *M. smegmatis* under hypoxia using a variation of the Wayne model (55), showing OD stabilization at 18-24 hours. Oxygen depletion was assessed qualitatively by methylene blue discoloration.

### ppGpp does not contribute to mRNA stabilization in hypoxia or carbon starvation

Given recent reports that ppGpp could directly inhibit the enzymatic activity of PNPase (40, 41), we wondered whether mRNA stabilization as observed in carbon starvation and hypoxia is regulated by ppGpp in mycobacteria. We obtained a double mutant strain of *M. smegmatis* (56) that lacks both genes implicated in the production of ppGpp (Δ*rel* Δ*sas2*), and compared the mRNA half-lives of a subset of genes to those of wild type mc^2^155 under hypoxia, log phase normoxia, and carbon starvation conditions. The Δ*rel* Δ*sas2* strain had a modest growth defect during adaptation to hypoxia and carbon starvation (Fig. 2A and 2C), as predicted (57). However, we found no significant decrease in mRNA stabilization in the mutant strain (Fig. 2B and 2D), indicating that the mRNA stabilization we observed under hypoxia and carbon starvation is independent from the stringent response. Interestingly, the mutant strain displayed increased mRNA stabilization for a few transcripts under carbon starvation conditions, which could be an indirect consequence of altered transcription rates (see discussion).

**Figure 2.**
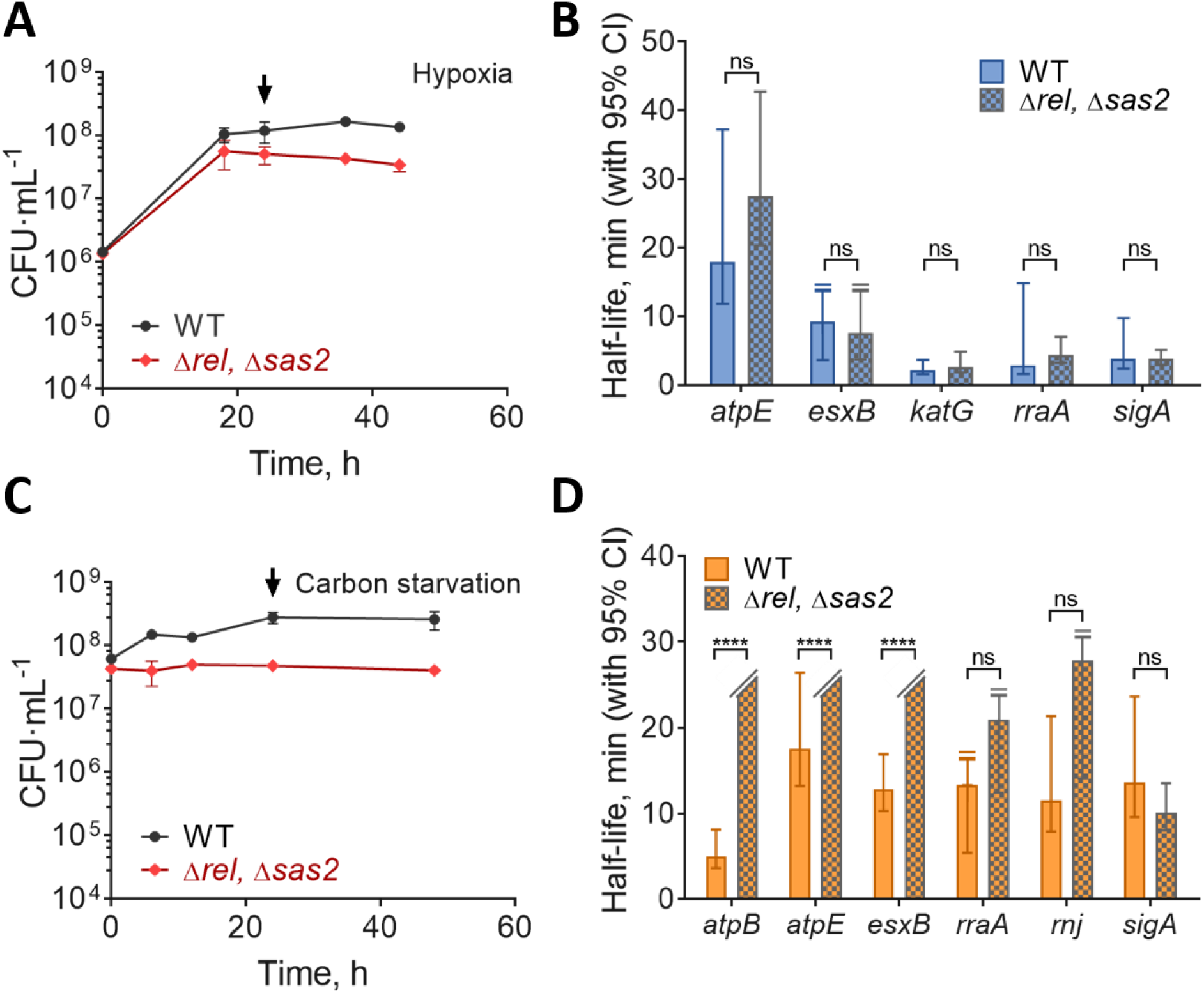
Transcript stabilization in hypoxia and carbon starvation is not dependent on the stringent response. (A) Growth kinetics for *M. smegmatis* mc^2^155 (WT) and Δ*rel*, Δ*sas2* strains cultured in 7H9 in flasks sealed at time 0. (B) Transcript half-lives for a set of genes 24 hours after sealing the hypoxia bottles (arrow in A). RNA samples were collected after blocking transcription with 150 μg·mL^−1^ RIF. (C) Bacteria were grown to log phase in 7H9 supplemented with ADC, glycerol, and Tween 80, then transferred to 7H9 supplemented with Tyloxapol only at time 0. (D) Transcript stability for a set of genes 22 hours after transfer to carbon starvation media (arrow in C). In A and C, the mean and SD of triplicate cultures is shown. In B and D, half-lives were compared using linear regression analysis (*n*=3). Error bars: 95% CI. **** *p*<0.0001, n.s. *p*>0.05. In cases where no degradation was observed or when the upper 95% CI limit was unbounded, the bar or upper error bar were clipped, respectively.

### Hypoxia-induced mRNA stability is reversible and independent of mRNA abundance

We wondered if the observed stress-induced transcript stabilization could be easily reversed by restoration of a favorable growth environment. To test this, we prepared 18 h hypoxia cultures, then opened the vials and agitated them for 2 min to re-expose the bacteria to oxygen before blocking transcription with RIF and sampling thereafter (Fig. 3A, top panel). We found that, for all transcripts tested, half-lives were significantly decreased compared to those observed under hypoxia and similar to those observed in log phase normoxia (Fig. 3B). While the mechanisms of stress-induced mRNA stabilization are largely unknown, multiple studies have reported inverse correlations between mRNA abundance and half-life in bacteria (3, 8, 51, 52). mRNA abundance is decreased for most transcripts tested in hypoxia-adapted *M. smegmatis*. We therefore considered the possibility that the dramatic increase in mRNA degradation upon re-exposure to oxygen was triggered by a rapid burst of transcription. Indeed, we found increased expression levels for three of five genes tested after two minutes of re-aeration, showing that transcription is rapidly induced upon return to a favorable environment (Fig. 3C). To test the idea that mRNA is destabilized by re-aeration as a consequence of a transcriptional burst and/or increased mRNA abundance, we modified our re-aeration experiment by blocking transcription with RIF one minute prior to re-exposure to oxygen (Fig. 3A, bottom panel). Surprisingly, every transcript tested was destabilized by the presence of oxygen despite the absence of new transcription. For most transcripts, the re-aeration half-lives were indistinguishable regardless of whether RIF was added prior to opening the vials or two minutes after (Fig. 3B). Our results therefore do not support the idea that changes in mRNA abundance alone can explain the mRNA stabilization and destabilization observed in response to changes in energy status.

**Figure 3.**
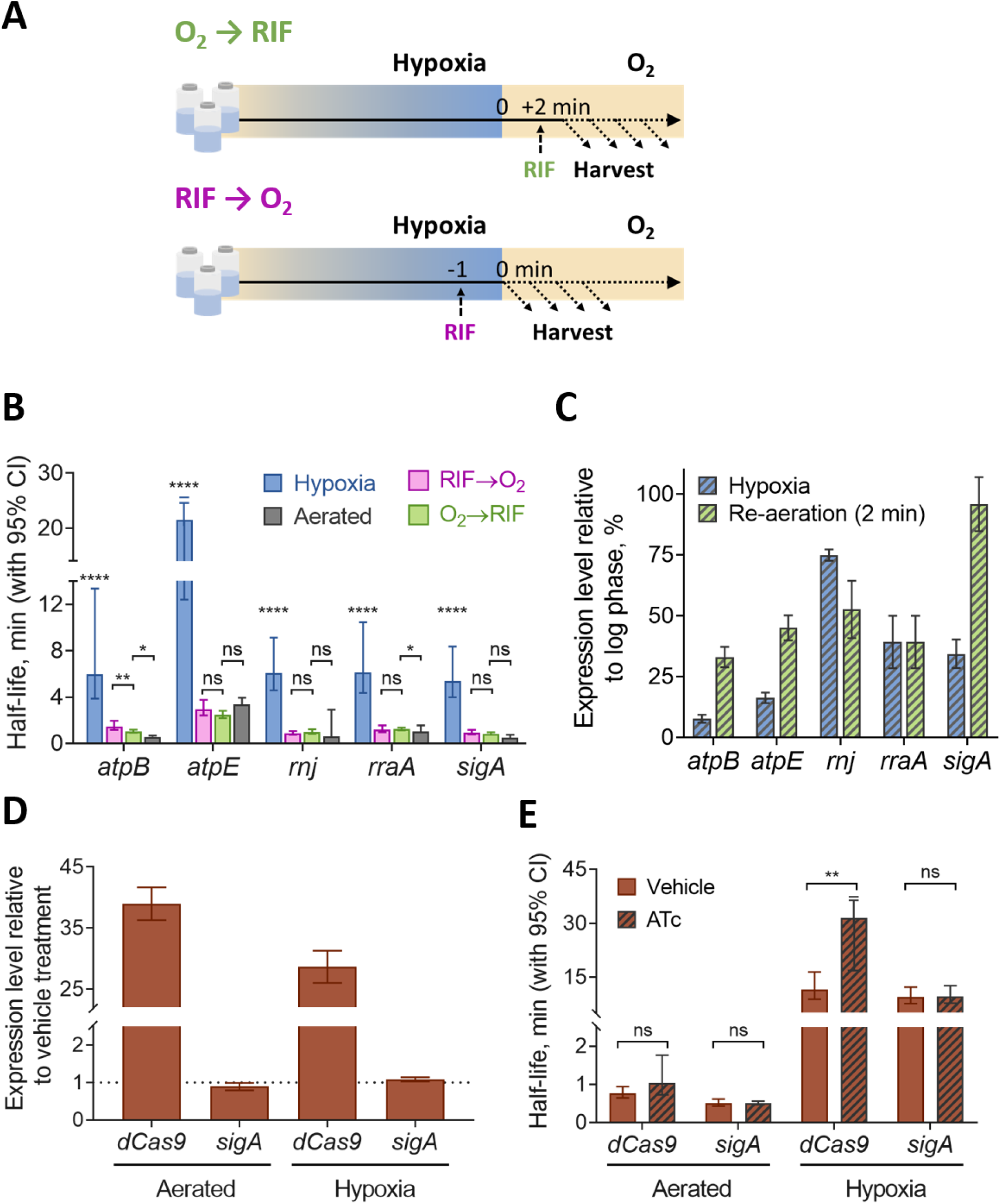
Hypoxia-induced mRNA stability is reversible and independent of mRNA abundance. (A) *M. smegmatis* was sealed in vials for 18 hours to produce a hypoxic environment, then re-exposed to oxygen for two minutes before transcription was inhibited RIF (top) or injected with RIF one minute prior to opening the vials and re-exposing to oxygen (bottom). (B) Transcript half-lives for a set of genes are displayed for log phase normoxia cultures, hypoxia (18 h), and re-aeration with RIF added either before or after opening the vials. Half-lives were compared by linear regression analysis (*n*=3). (C) Expression levels of transcripts in hypoxia (18 h) or 2 min re-aeration relative to the expression levels in log phase normoxia cultures (percentage). Error bars: SD. (D) Expression levels of transcripts in hypoxia (18 h) or log phase normoxia after being treated with 200 ng·mL^−1^ ATc for 1 h or 10 min, respectively, to induce *dCas9* overexpression, relative to the expression levels in a H_2_O vehicle treatment (percentage). Error bars: SD. (E) Transcript half-lives for *dCas9* and *sigA* for log phase normoxia and hypoxia (18 h) after induction of *dCas9* with ATc or vehicle treatment as shown in D. In B and E, degradation rates were compared using linear regression (*n*=3), and half-lives were determined by the reciprocal of the best-fit slope. Error bars: 95% CI. * *p*<0.05, ** *p*<0.01, **** *p*<0.0001, n.s. *p*>0.05.

We wanted to further explore if mRNA abundance alone could influence transcript degradation. Hence, we obtained a *M. smegmatis* strain encoding *dCas9* and a non-specific sgRNA under control of an ATc-inducible promoter (58) and compared the *dCas9* transcript stability under hypoxia and log phase normoxic conditions after ATc induction or at basal levels of expression. Our results showed that despite a 34-fold transcript upregulation following ATc induction, the half-life of *dCas9* mRNA was not significantly different from the no-drug control under log phase normoxia. Under hypoxia, its 28-fold upregulation was associated with a modest increase in *dCas9* mRNA half-life when compared to the no-drug control (Fig. 3D and 3E). Taken together, our results show that increased mRNA abundance does not necessarily result in a faster decay rate.

### mRNA stability is modulated independently of RNase protein levels

Another potential explanation for increased mRNA degradation after re-aeration is the up-regulation of mRNA degradation proteins such as RNase E. To assess the role of a sudden burst in protein levels we used two approaches. First, we constructed *M. smegmatis* strains encoding FLAG-tagged RNase E, cMyc-tagged PNPase, or cMyc-tagged predicted RNA helicase msmeg_1930. PNPase is an essential exoribonuclease. We determined protein levels by western blotting during log phase, in 18 h hypoxia, and after 18 h hypoxia followed by 2 min re-aeration. As shown in Fig. 4A, levels of all three of these predicted RNA degradation proteins remained unchanged in the three conditions.

**Figure 4.**
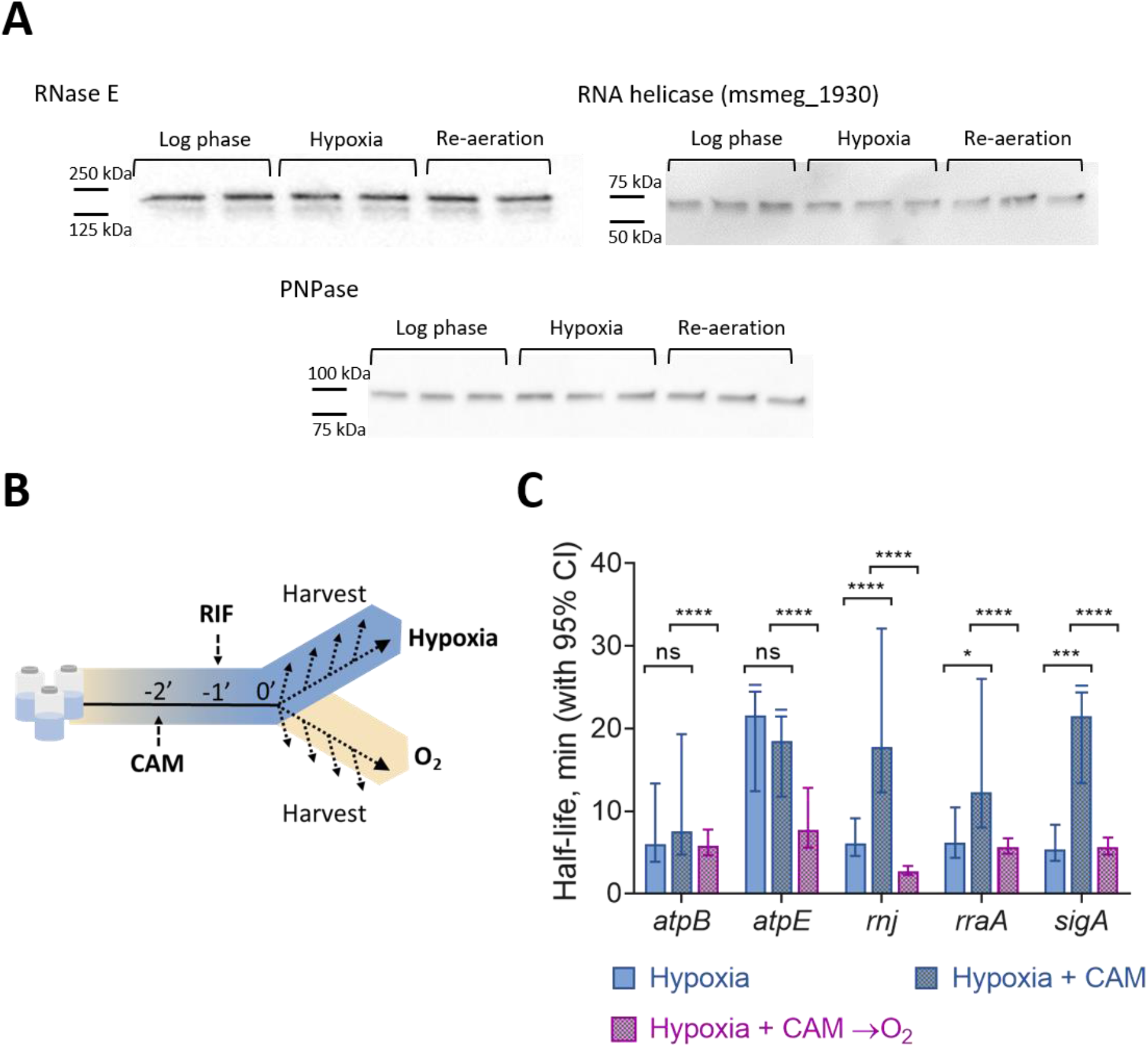
mRNA stability is regulated independently of degradation protein levels. (A) Western blotting for FLAG-tagged RNase E, and c-Myc-tagged PNPase or RNA helicase (msmeg_1930) in *M. smegmatis* in log phase normoxia, hypoxia (18 h), and 2 min re-aeration. Samples were normalized to total protein level, which were similar on a per-OD basis in all conditions. (B) Translation was inhibited in hypoxic cultures by 150 μg·mL^−1^ CAM 1 min before adding 150 μg·mL^−1^ RIF. RNA was harvested at time points beginning 2 min after adding CAM. (C) Transcript half-lives for samples from hypoxic cultures with the drug vehicle (ethanol), hypoxic cultures after translation inhibition, and 2 min re-aeration after translation inhibition. Degradation rates were compared using linear regression (*n*=3), and half-lives were determined by the reciprocal of the best-fit slope. Error bars: 95% CI. n.s., *p*>0.05, * *p*<0.05, *** *p*<0.001, **** *p*<0.0001.

Because we do not know all of the proteins that contribute to mRNA degradation in mycobacteria, our second approach was to test the global importance of translation in re-aeration-induced mRNA destabilization. We blocked translation with chloramphenicol (CAM) in 18 h hypoxia cultures and then added RIF. Samples were collected for cultures that remained under hypoxia as well as those that were re-exposed to oxygen for 2 min (Fig. 4B). Our results showed that there is destabilization of mRNA after re-aeration even in the absence of protein synthesis (Fig. 4C), though not to the extent we observed in Fig. 3B. These results suggest that re-aeration-induced destabilization does not require synthesis of new RNA degradation proteins. The mRNA stabilization induced by CAM itself is likely related to its mechanism of action. CAM inhibits elongation by blocking the 50S ribosomal subunit from binding tRNAs, preventing peptidyl transferase activity (59–61) and causing ribosomal stalling (62). Thus, consistent with our data, others have reported global stabilization of mRNA pools when elongation inhibitors, but not initiation inhibitors, are used for example in log phase cultures of *E. coli* (62) or in yeast (63). We hypothesize that stalled ribosomes may increase mRNA stability by masking RNase cleavage sites. However, we observe mRNA destabilization in response to re-aeration despite this effect (Fig. 4C). Taken together, our data suggest that tuning of protein levels is not the primary explanation for mRNA stabilization during early adaptation to hypoxia.

### mRNA stability is modulated in response to changes in metabolic status

The rapidity of mRNA destabilization in response to re-aeration suggested that mRNA degradation is tightly regulated in response to changes in energy metabolism. We tested this hypothesis by treating log phase cultures of *M. smegmatis* with 5 μg·mL^−1^ bedaquiline (BDQ), a potent inhibitor of the ATP synthase F0F1 (64). We used minimal media that contained acetate as the only carbon source (MMA) in order to make the respiratory chain the sole source of ATP synthesis. After 30 min exposure, intracellular ATP levels were reduced by more than 90% in BDQ-treated cells, when compared to cells treated with the drug vehicle (DMSO), without affecting cell viability (Fig. 5A and 5B). We then measured half-lives for a set of transcripts under these conditions. mRNA half-lives were dramatically increased in BDQ-treated cells for most of the genes we tested (Fig. 5C), indicating that mRNA degradation rates are rapidly altered in response to changes in energy metabolism status.

**FIG 5.**
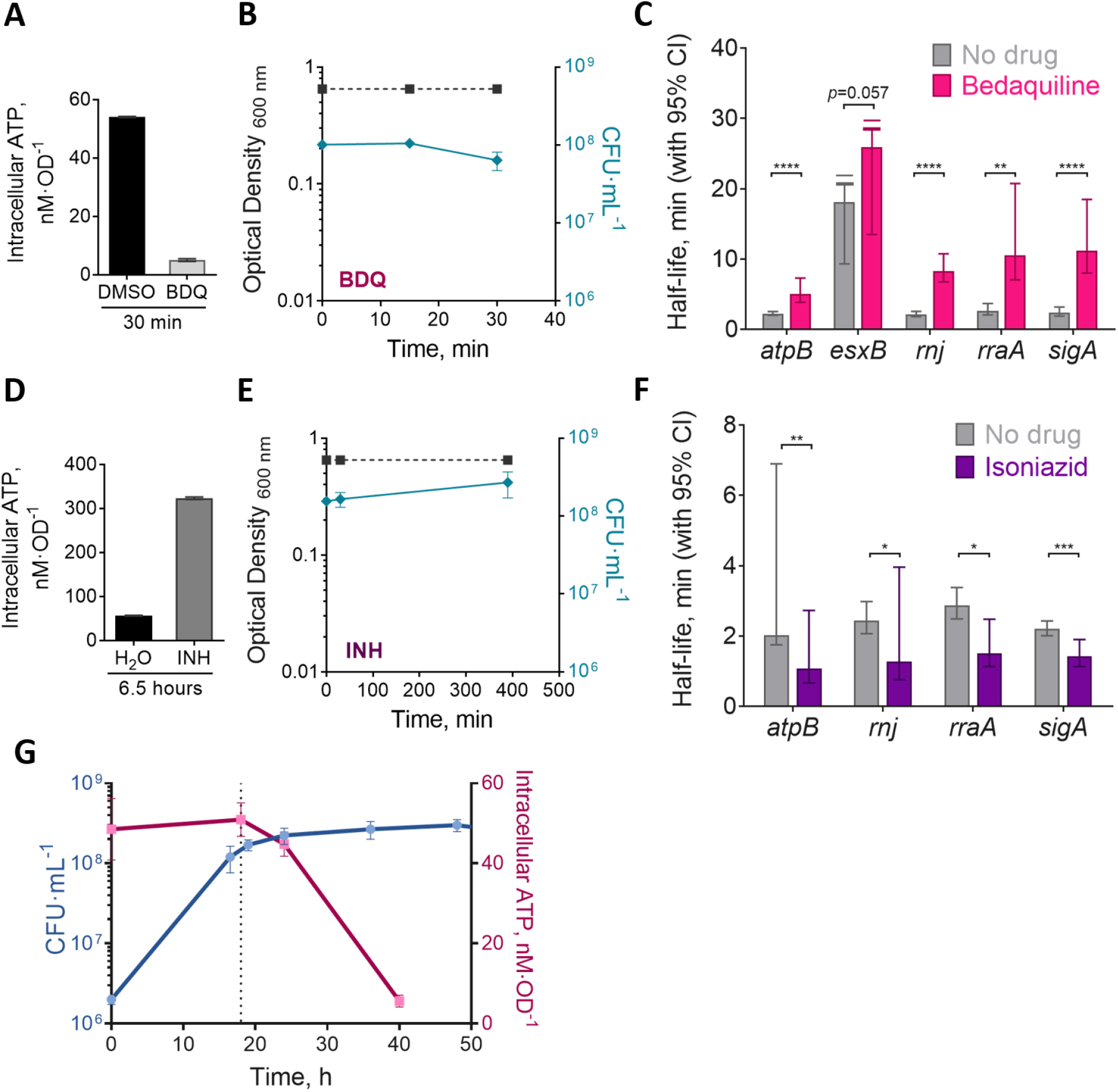
mRNA stability is modulated in response to changes in metabolic status. (A) *M. smegmatis* was cultured in MMA media for 22 hours to OD_600_ 0.8 before being treated with 5 μg·mL^−1^ BDQ or the vehicle (DMSO) for 30 min. Intracellular ATP was determined using the BacTiter-Glo kit. (B) Growth kinetics for *M. smegmatis* from panel A in presence of BDQ. (C) Transcript half-lives for a sub-set of transcripts collected during intracellular ATP depletion (30 min with BDQ) or at the basal levels (30 min with DMSO). (D) As in panel A, but for *M. smegmatis* treated with 500 μg·mL^−1^ INH or the vehicle (H_2_O) for 6.5 hours. (E) Growth kinetics for *M. smegmatis* from panel D in presence of INH. (F) Transcript half-lives for a sub-set of transcripts after 6.5 h of INH or vehicle treatment. (G) Growth kinetics for *M. smegmatis* transitioning into hypoxia, and intracellular ATP levels at different stages. The dotted line represents the time at which transcript stability analysis were made for the hypoxia (18 h) condition for Figures 1–4. In C and F, half-lives were compared using linear regression analysis (*n*=3). Error bars: 95% CI. * *p*<0.05, ** *p*<0.01, *** *p*<0.001, **** *p*<0.0001.

We then wondered if we could increase mRNA degradation rates by increasing intracellular ATP levels. To test this, we treated *M. smegmatis* cultures with isoniazid (INH) a pro-drug that interferes with the synthesis of mycolic acids, but that also leads to an accumulation of intracellular ATP due to increased oxidative phosphorylation (65). We exposed *M. smegmatis* to 500 μg·mL^−1^ INH for 6.5 hours to confirm that we had achieved bacteriostasis (the *M. smegmatis* doubling time in MMA media is approximately six hours). As shown in Fig. 5D, INH caused a dramatic increase in intracellular ATP after 6.5 h without affecting cell viability (Fig. 5E). Remarkably, mRNA half-lives were significantly decreased in response to INH (Fig. 5F). To our knowledge, this is the first report of bacterial mRNA being destabilized rather than stabilized in response to a growth-impairing stressor. Our results indicate that mRNA stability is regulated not in response to growth status per se, but rather to energy metabolism. Although we interpreted ATP levels as a reflection of metabolic status in our INH and BDQ assays, the coupling between mRNA degradation and metabolic status does not appear to be mediated by ATP directly. We measured ATP levels in *M. smegmatis* cultures during the transition to hypoxia-induced growth arrest, and found that although ATP levels ultimately decrease in hypoxia as has been reported elsewhere (66, 67), mRNA stabilization precedes the drop in ATP levels (Fig. 5G).

## DISCUSSION

Many stressors cause bacteria to slow or stop growth, and this is usually associated with increased mRNA stability (3–9, 11–13). Many of these same stressors reduce energy availability (66, 67), requiring reductions in energy consumption and optimization of resource allocation. We speculate that the decreased mRNA turnover that accompanies such conditions may be an energy conservation mechanism. For Mtb, hypoxia can lead to generation of bacterial subpopulations with varying degrees of antibiotic tolerance (68–70), facilitating bacterial survival and the acquisition of drug resistance-conferring mutations. Understanding the mechanisms that support the transitions into non-growing states, and subsequent survival in these states, is therefore of great importance.

The transcriptome of Mtb has been previously shown to be stabilized under cold shock and hypoxia (3). Here, we found that *M. smegmatis* also dramatically stabilized its mRNA in response to carbon starvation and oxygen depletion. For the first time, to our knowledge, we tested the speed at which this stabilization is reversed in mycobacteria upon restoration of energy availability. Remarkably, mRNAs are rapidly destabilized within minutes of re-aeration of hypoxic cultures, suggesting that tuning of mRNA degradation rates is an early step in the response to changing energy environments.

The most straightforward explanation for stress-induced mRNA stabilization would seem to be downregulation of the mRNA degradation machinery. Indeed, under hypoxic conditions, RNase E is downregulated at the transcript level, and abundance of cleaved RNAs is notably reduced (71). However, we found that protein levels were unchanged for three proteins predicted to be core components of the mRNA degradation machinery. This is largely consistent with what was reported for Mtb in a quantitative proteomics study (37), although in that case there was an apparent reduction in levels of a predicted RNA helicase. To address this question in a more agnostic fashion, we tested the importance of translation for transcript destabilization upon re-exposure of hypoxic cultures to oxygen. However, re-aeration triggered increased transcript degradation even in the absence of new protein synthesis. Regulation of degradation protein levels therefore does not appear to contribute to mRNA stabilization during the initial response to energy stress. However, we found that upon longer periods of hypoxia, transcripts were stabilized to a greater extent than what we observed 18 hours after sealing the vials. This suggests that mRNA stabilization progressively increases, and may be the product of multiple mechanisms. As this work focused on the initial transition into hypoxia-induced growth arrest, we cannot discount the possibility that downregulation of the RNA degradation machinery is important for further mRNA stabilization in later hypoxia stages.

Interestingly, we found greater mRNA stabilization in hypoxic cultures treated with CAM. This may result from stalled ribosomes (59, 61) masking RNase cleavage sites. Furthermore, the burst of transcription upon re-aeration is blocked by the presence of CAM, causing up to a four-fold decrease in transcript abundance in the CAM treated cultures when compared to the vehicle treated cultures. This is consistent with the idea that transcription and translation are physically coupled, and blocking translation therefore prevents RNA polymerase from efficiently carrying out transcript elongation, as has been reported for *E. coli* (72–76).

Transcript abundance has been found to be inversely correlated with mRNA stability in exponentially growing bacteria (3, 8, 51, 52, 77), and experimental manipulation of transcription rates of subsets of genes resulted in altered degradation rates (3, 52). Together, these studies suggest that high rates of transcription inherently increase degradation rates. We report here that during oxygen depletion transcript levels are reduced in *M. smegmatis*, which led us to ask if increased transcript half-lives under stress are a direct result of reduced mRNA levels. However, our data are inconsistent with this idea; mRNA is rapidly destabilized upon re-aeration even in the absence of new transcription. We note that one study reported a weak positive correlation between mRNA abundance and stability in log phase *E. coli* (12), while another reported mRNA abundance to be positively correlated with stability in carbon-starved *Lactococcus lactis* (8). Taken together, these observations and our own suggest that the relationship between mRNA stability and abundance is not yet fully understood and may be fundamentally different in growth-arrested bacteria.

The rapid reversibility of hypoxia-induced mRNA stabilization suggests that mRNA decay and energy metabolic status are closely linked. Consistent with this idea, we have shown that drug-induced energy stress causes mRNAs to be stabilized, while mRNA decay is increased by a drug that induces a hyperactive metabolic state. To our knowledge this is the first demonstration that the rate of bacterial mRNA degradation can be decoupled from growth rate, and suggests that mRNA decay is controlled by energy status rather than growth rate per se. The mechanism by which energy status and mRNA decay are coupled remains elusive; the stringent response is not required, and the stabilization of mRNA during adaptation to hypoxia precedes a decrease in ATP levels. Possible explanations that should be investigated in future work include ribosome occupancy, the presence of other RNA-binding proteins, and the subcellular localization of mRNAs and the RNA degradation machinery.

## METHODS

### Strain and culture conditions

*Mycobacterium smegmatis* strain mc^2^155 or derivatives (Table 1) were grown in *rich medium*, Middlebrook 7H9 supplemented with ADC (Albumin Dextrose Catalase, final concentrations 5 g·L^−1^ bovine serum albumin fraction V, 2 g·L^−1^ dextrose, 0.85 g·L^−1^ NaCl, and 3 mg·L^−1^ catalase), 0.2% glycerol and 0.05% Tween 80 at 200 rpm and 37°C to an OD_600_ of ~0.8, unless specified otherwise. For the hypoxic cultures, we used a modification of the Wayne model (55). Briefly, *M. smegmatis* was cultured in 30.5 × 58 mm serum bottles (Wheaton, 223687, 20 mL) using *rich medium* and an initial OD_600_=0.01. The bottles were sealed with a vial crimper (Wheaton, W225303) using rubber stoppers (Wheaton, W224100-181) and aluminum seals (Wheaton, 224193-01). The cultures were grown at 37 °C and 200 rpm to generate hypoxic conditions. Oxygen levels were qualitatively monitored using methylene blue.

**TABLE 1.**
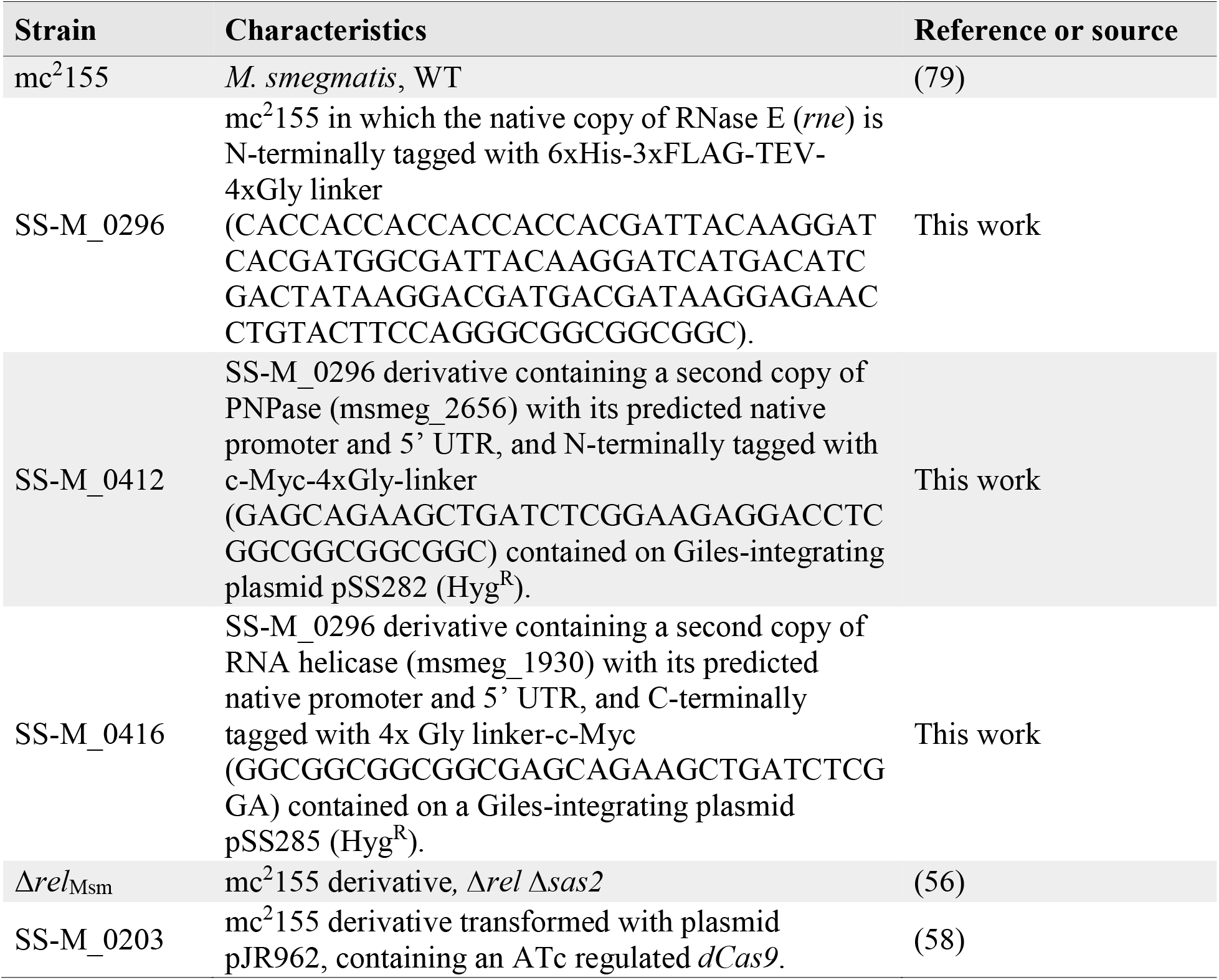
Strains used and sources

Carbon starvation cultures were prepared using log phase cells (OD_600_=0.8) grown in *rich medium*. Cultures were rinsed with *carbon starvation medium* (Middlebrook 7H9 supplemented with 5 g·L^−1^ bovine serum albumin fraction V, 0.85 g·L^−1^ NaCl, 3 mg·L^−1^ catalase and 0.05 % Tyloxapol) and centrifuged for 5 min at 3,214 × g at 4°C. After three rinses, the pelleted cells were resuspended in *carbon starvation medium* to an OD_600_= 0.8 and incubated at 200 rpm and 37°C.

### RNA extraction and determination of mRNA stability

Biological triplicate cultures were treated with rifampicin (RIF) to a final concentration of 150 μg·mL^−1^ to halt transcription and RNA was extracted at various time points thereafter. For exponentially growing cells in normoxia and cells in carbon starvation, 7 mL samples (OD_600_ ~0.8) were collected per replicate and time point after blocking transcription. Samples and were snap-frozen in liquid nitrogen. For hypoxic samples, degassed RIF was injected using a 30G needle, and all samples were sacrificially collected per time point and replicate (7 mL, OD_600_ ~0.8) and snap-frozen in liquid nitrogen within 6 seconds of unsealing the bottles. Time points were taken at different intervals after adding RIF depending on the experiment.

Samples were stored at −80°C and thawed on ice immediately before RNA extraction. Cells were centrifuged for 5 min at 3,214 × g at 4°C, and supernatants removed completely. Working on ice, the pellet was resuspended in 1 mL of TRIzol (Invitrogen), transferred to 2 mL disruption tubes (OPS Diagnostics 100 μm zirconium lysing matrix, molecular grade) for cell lysis using a FastPrep-24 5G (MP) with 3 cycles of 7 m·s^−1^ for 30 s, with 2 min on ice after each cycle. 300 μL chloroform was added to each sample, mixed and centrifuged for 15 min at 21,130 × g and 4 °C. RNA was recovered from the aqueous layer and purified after DNase digestion in-column using the Direct-zol RNA MiniPrep kit according to the manufacturer’s instructions. A NanoDrop 2000c (Thermo) was used to determine RNA concentrations and 1% agarose gels were used to verify RNA integrity.

For cDNA synthesis, 600 ng of total RNA were mixed with 0.83 μL 100 mM tris pH 7.5 and 0.17 μL 3 mg·mL^−1^ Random Primers (NEB) to a volume of 5.25 μL, denatured at 70°C for 10 min and snap-cooling for 5 min. Reverse transcription was performed for 5 hours at 42 °C using 100 U of ProtoScript^®^ II Reverse Transcriptase (NEB), 10 U of RNase Inhibitor (Murine, NEB), 0.5 mM each dNTP mix and 5 mM DTT in a final volume of 10 μL. RNA was degraded at 65°C for 15 min by addition of 5 μL each 0.5 mM EDTA and 1 N NaOH, halting the reaction with 12.5 μL of 1 M Tris-HCl pH 7.5. cDNA was purified using the MinElute PCR Purification Kit (Qiagen) according to the manufacturer instructions. mRNA abundance (A) over time (*t*) was determined for different genes (primers are listed in Table 2) by quantitative PCR (qPCR) using iTaq SYBR Green (Bio-Rad) with 400 pg of cDNA and 0.25 μM each primer in 10 μL reactions, with 40 cycles of 15 s at 95°C followed by 1 min at 61°C (Applied Biosystems™ 7500 Real-Time PCR). Abundance was expressed as the -Ct (or the log_2_A(*t*)). Linear regression was performed on -Ct values versus time where the negative reciprocal of the best-fit slope estimates mRNA half-life. In many cases the decay curves were biphasic, where a rapid period of decay was followed by a period of slow or undetectable decay. In these cases, only the initial, steeper slope was used for calculation of half-lives.

**TABLE 2.**
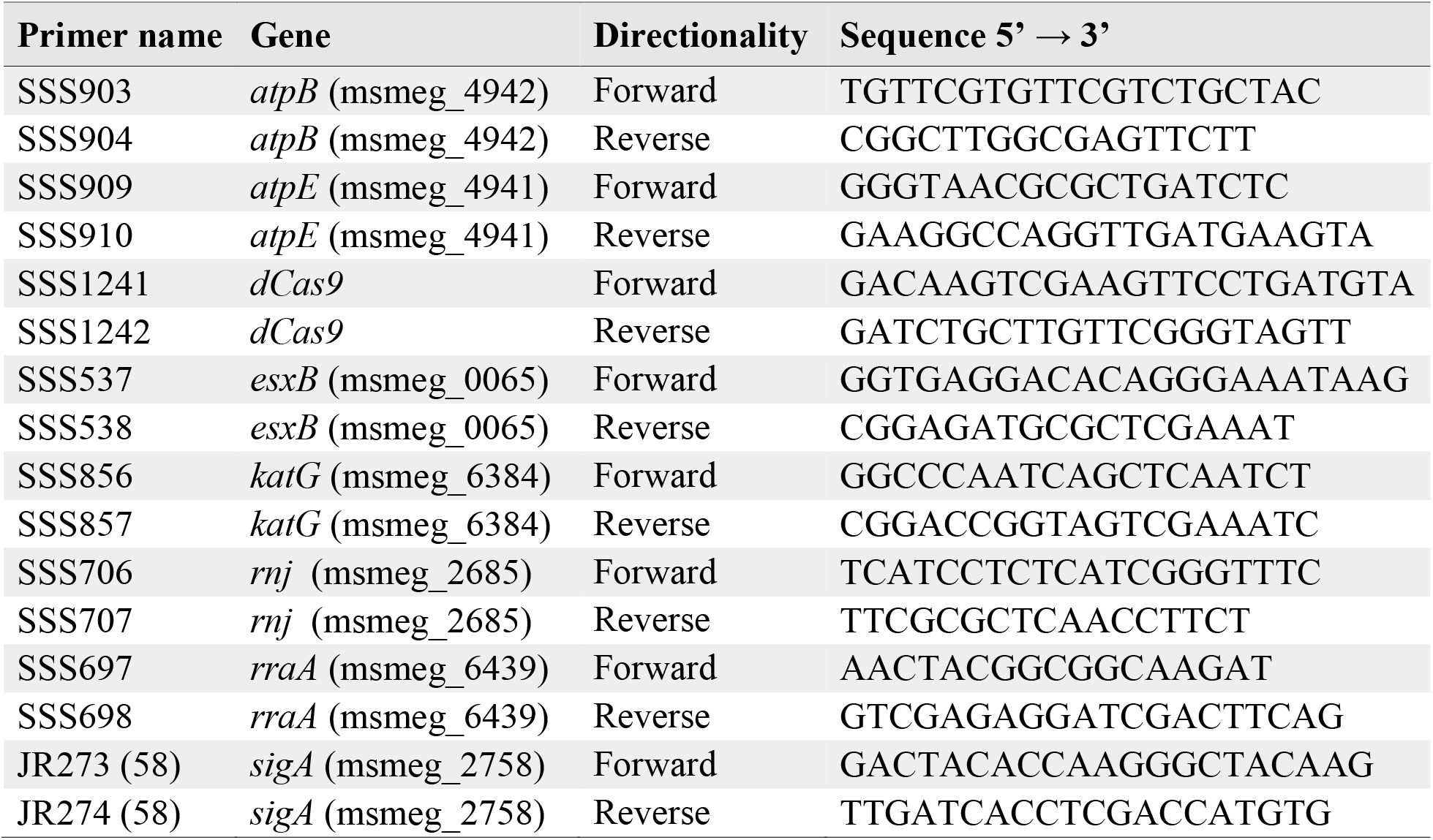
Primers for qPCR

### mRNA stability measured during re-aeration and translational inhibition

Translation was halted in normoxia and hypoxia cultures by chloramphenicol at a final concentration of 150 μg·mL^−1^. 1 min after adding chloramphenicol, rifampicin was added, and time point samples were collected starting 1 min afterwards.

Re-aeration experiments were done using hypoxia cultures 18 hours after the sealing. Briefly, each bottle was opened and the contents transferred to a 50 mL conical tube. Rifampicin was added either 1 min before (transcription inhibition during hypoxia) or 2 min after opening the bottles (transcription inhibition after re-aeration). Samples were taken 2, 7, 12, 17, and 32 min after opening the bottles and snap-frozen in liquid nitrogen as described before. Samples were collected in triplicate, steps prior to freezing were performed at 37°C, and incubation of the samples in either container was done at 200 rpm.

### Construction of 6xHis-3xFLAG tagged chromosomal RNase E *M. smegmatis* strain

The RNase E-tagged strain (SS-M_0296) was built using a two-step process. Plasmid pSS250 was derived from pJM1 (78) and contained 1 kb of the sequence upstream and downstream of the *rne* (msmeg_4626) start codon, with the sequence coding for the 6xHis-3xFLAG-TEV-4xGly linker inserted after the start codon of *rne*. Constructs were built using NEBuilder HiFi DNA Assembly Master Mix kit (E2621). Integrants were selected based on Hyg^R^ (200 μg·mL^−1^ hygromycin) and confirmed by sequencing. Counter-selection with 15% sucrose was followed by PCR screening to identify isolates that subsequently underwent second crossovers resulting in loss of the plasmid and retention of the *rne*-FLAG-tagged sequence.

### Construction of c-Myc tagged Helicase and PNPase strains

The PNPase-tagged strain (SS-M_0412) was built by inserting a second copy of *pnp* (msmeg_2656) with an N-terminal c-Myc-4xGly-linker along with its predicted native promoter and 5’ UTR at the Giles phage integration site (plasmid pSS282) into strain SS-M_0296. The RNA helicase-tagged strain (SS-M_0416) was constructed in a similar way but using a C-terminal 4xGly-linker-c-Myc tag on msmeg_1930 (plasmid pSS285).

### Alteration of intracellular ATP with bedaquiline and isoniazid

Cells were grown as described before to an OD_600_ of ~1.0, rinsed two times in *Minimal Media Acetate wash* (final concentrations are 0.5 g·L^−1^ L-asparagine, 1 g·L^−1^ KH_2_PO_4_, 2.5 g·L^−1^ Na_2_HPO_4_, 0.5 g·L^−1^ MgSO_4_•7H_2_O, 0.5 mg·L^−1^ CaCl_2_, 0.1 mg·L^−1^ ZnSO_4_, 0.1% CH_3_COONa, 0.05% tyloxapol, pH 7.5) using 5 min centrifugation steps at 3,214 × g and 4°C, and finally resuspended in Minimal Media Acetate with ferric ammonium citrate (MMA, *Minimal Media Acetate wash* + 50 mg·L^−1^ ferric ammonium citrate) at an OD_600_=0.07. The cells were then grown for 24 hours at 37°C and 200 rpm to an OD_600_ of ~0.8. To remove the high amounts of extracellular ATP, 30 minutes before drug treatment the cells were rinsed in pre-warmed *Minimal Media Acetate wash* as described before, resuspended in pre-warmed MMA, and returned to the incubator. Either bedaquiline (BDQ), isoniazid (INH) or their vehicles were added to the cell cultures to a final concentration of 5 μg·mL^−1^ or 500 μg·mL^−1^, respectively. Cultures were incubated as described before, and samples were taken 30 min after adding BDQ, or 6.5 h after adding INH for half-life-estimation and ATP-determination assays.

For half-life measurements, samples for BDQ were taken 0, 3, 6, 9, 12, 15 and 21 min after adding RIF. Samples for INH were taken 0, 4, 8 and 12 min after adding RIF. All samples were collected in triplicates. RNA extraction for cultures in MMA was performed as indicated before with the following modifications: cell disruption was performed using 2 mL tubes prefilled with Lysing Matrix B (MP) and 3 cycles of 10 m·s^−1^ for 40 s; RNA was recovered from the aqueous layer by isopropanol precipitation and resuspension in RNase-free H_2_O; samples were treated with 5U of TURBO™ DNase (Ambion) in presence of 80 U of RNase Inhibitor, Murine (NEB) for 1 hour at 37°C and under agitation. RNA was purification with an RNeasy Mini Kit (Qiagen) according to the manufacturer’s specifications.

### Intracellular ATP estimation

ATP was estimated using the BacTiter-Glo kit (Promega). For BDQ or INH treatments in MMA, after the treatment periods stated above, 1 mL of culture was pelleted at ~21°C for 1 min at 21,130 x g. The supernatant was removed and the cells were resuspended in 1 mL of pre-warmed MMA containing either BDQ, INH or the vehicle to match the prior treatment condition. Immediately after, 20 μL samples were transferred to a white 384-well plate (Greiner bio-one) containing 80 μL of the BacTiter-Glo reagent and mixed for 5 minutes at room temperature. Luminescence was measured in a VICTOR^3^ plate reader (PerkinElmer) (intracellular ATP). We included controls for the supernatant collected (extracellular ATP), media + drug/vehicle (background), and freshly made ATP standards for each reading.

To estimate the intracellular ATP in normoxia and hypoxia Middlebrook 7H9 cultures, 20 μL samples were collected at 37°C and immediately combined with the reagent to measure total ATP (intracellular + extracellular). From the same cultures, 1 mL samples were syringe-filtered (PES 0.2 μm) and the filtrate combined with the reagent to measure extracellular ATP. Luminescence was measured as described above. Intracellular ATP was calculated by subtracting the extracellular ATP values from the total ATP values. Hypoxia samples were sacrificially harvested per time point/replicate and combined with the reagent in <6 seconds. The respective controls and ATP standards were also included for each reading. All samples were measured in biological triplicate, and in at least two independent experiments.

## AUTHOR CONTRIBUTIONS

DVB and SSS conceived and design the experiments. DVB and YZ performed the experiments. LGZ performed part of the experiments in Fig. 3. DVB, TA and SSS analyzed the data. DVB and SSS wrote the manuscript.

## ACKNOWLEDGEMENTS

This work was supported by NSF CAREER award 1652756 to SSS. DVB was partially supported by the LASPAU Fulbright Foreign Student Program. We thank all members of the Shell lab for technical assistance and helpful discussions. The thank Dr. Christina Stallings, Dr. Jeremy Rock and Dr. Sarah Fortune for generously providing strains.

